# Environmental DNA reveals long-term persistence of a *Midichloria*-like bacterium in a rainbow trout aquaculture and links *Ichthyopthirius multifiliis* with the red mark syndrome

**DOI:** 10.64898/2026.04.16.718929

**Authors:** Davide Vecchio, Ylenia Siviglia, Alessandro Allievi, Elisa Fesce, Paolo Losi, Carlo Croci, Leandro Gammuto, Luca Ilahiane, Sophie Melis, Alessandra Cafiso, Nicola Ferrari, Giulio Petroni, Valentina Serra, Perla Tedesco, Michele Castelli

## Abstract

Red Mark Syndrome (RMS) is a widespread skin disease affecting rainbow trout (*Onchorhynchus mykiss*). It provokes substantial economic losses in aquaculture, and is putatively caused by a *Rickettsiales* bacterium named *Midichloria*-like organism (RMS-MLO), which is strongly associated with RMS lesions. However, RMS-MLO ecology and epidemiology in aquaculture systems remain poorly understood. In this study, we analysed environmental DNA to monitor the presence of RMS-MLO and its putative vector *Ichthyophthirius multifiliis* in a trout farm in Northern Italy over one year. Water and sediment samples were monthly collected from multiple water tanks. RMS-MLO was consistently detected by PCR throughout the study in all trout-containing tanks, both in water and sediment samples, but never in the trout-free inflow tank. We did not observe an increase in RMS-MLO abundance during the single RMS outbreak recorded nor in relation with the co-occurrence of *I. multifiliis*. Our findings indicate a long-term persistence of RMS-MLO in the aquaculture, possibly as a consequence of infections with low prevalence or abundance, rather than its entry from the external environment at the time of RMS outbreaks. Additionally, hints were recorded for a potential role of free-living aquatic microeukaryotes as additional occasional reservoirs. In contrast, *I. multifiliis* was negatively related with RMS-MLO, while it significantly increased in abundance during the RMS outbreak, particularly in the inflow tank. This supports that, rather than a stable reservoir, *I. multifiliis* may act as a facilitator of RMS outbreaks, which might indeed be triggered by the entry of this parasite in trout farms.

## 1. Introduction

Aquaculture production has steadily increased in recent years, accounting for approximately 1% of total global merchandise trade value in 2023 (FAO, 2023). In aquaculture systems, high fish population density and stress can facilitate the emergence and spread of infectious diseases (Blandford et al., 2018; Murray and Peeler, 2005), which represent a major source of economic losses (Lafferty et al., 2015; Meyer, 1991).

The rainbow trout *Oncorhynchus mykiss* is one of the most widely farmed fishes worldwide (D’Agaro et al., 2022). It can be affected by several diseases of bacterial and mycotic origin, such as furunculosis, columnaris disease, and saprolegniosis (Oidtmann et al., 2013). Among those, the Red Mark Syndrome (RMS) was first described in 2003 from UK cases (Verner-Jeffreys et al., 2008), although it had previously been reported as “strawberry disease” in the US (Metselaar et al., 2022; Oidtmann et al., 2013; Schmidt-Posthaus et al., 2009). RMS is characterised by single or multiple bright red skin lesions, most commonly located on the flanks, and is typically accompanied by pronounced inflammation and subcutaneous oedema (Galeotti et al., 2023; McCarthy et al., 2013; Metselaar et al., 2022). It is classified into three severity types based on lesion size and colour, presence of exudate, scale loss, and tissue erosion (M. Galeotti et al., 2021). Histologically, the disease is characterised by a progressive leukocyte infiltration, with notable features observed in severe lesions including panniculitis, myositis extending to the myosepta, and acanthosis (M. Galeotti et al., 2021). The disease onset and resolution are negatively correlated with water temperature, with highest incidence at around 12°C (Orioles et al., 2022a).

Although RMS is non-lethal (with one not officially confirmed exception (Oh et al., 2019)), its high morbidity coupled with downgraded affected fish at harvest causes significant economic losses (Metselaar et al., 2022). This syndrome has been reported worldwide, namely repeatedly across Europe (Galeotti et al., 2011, 2017b; Metselaar et al., 2022; Schmidt-Posthaus et al., 2009), in north and south America (Ortega et al., 2023; Sandoval et al., 2016), and in Asia (Metselaar et al., 2022; Oh et al., 2019). In nation-wide surveys conducted in Italy and Denmark, a significant proportion of trout farms (∼30%) was found affected (Orioles et al., 2023; Schmidt et al., 2018), while up to 20% of total fish in a single farm may be infected during an outbreak (Galeotti et al., 2017b; Marco Galeotti et al., 2021).

RMS is transmissible (Verner-Jeffreys et al., 2008; von Gersdorff Jørgensen et al., 2019a), with over 80% of naïve trout developing the disease within 60 days of cohabitation with RMS-infected individuals (Orioles et al., 2022a; Schmidt et al., 2021), and likely caused by a bacterium (Metselaar et al., 2010; Schmidt et al., 2021). Although the aetiological agent has not yet been formally demonstrated, multiple evidences strongly point to an uncharacterised *Midichloria*-related *Rickettsiales* bacterium, accordingly named RMS-MLO (*Midichloria*-like organism) (Cafiso et al., 2016). Indeed, RMS-MLO has been repeatedly detected by PCR in association with RMS skin lesions (Cafiso et al., 2016; Lloyd et al., 2011, 2008; Metselaar et al., 2020, 2010; Pardo et al., 2024; Zarantonello et al., 2025), although its abundance was not found to correlate with lesion severity (Cafiso et al., 2016; Galeotti et al., 2023). Ultrastructural investigations revealed an intracytoplasmic bacterium in RMS lesions, which may represent RMS-MLO (Galeotti et al., 2017a; Orioles et al., 2022a). Nevertheless, experimental investigations on RMS-MLO, including a formal test of involvement in RMS, are hampered by the infeasibility of its isolation in culture (Cafiso et al., 2016; Orioles et al., 2022b), which is a common feature among *Rickettsiales* bacteria, being intracellular and host-dependent for their replication (Salje, 2021).

Pathogenic *Rickettsiales* are typically transmitted by haematophagous arthropod vectors (Abdad et al., 2018; Dantas-Torres et al., 2012), while most aquatic *Rickettsiales* are hosted by unicellular eukaryotes (Castelli et al., 2024, 2016). Accordingly, a possible role in the transmission of RMS-MLO was hypothesised for the ciliate protozoan *Ichthyophthirius multifiliis*, a common skin parasite of rainbow trout and other fish species in aquaculture (Jørgensen, 2017; Matthews, 2005). Indeed, DNA from RMS-MLO (or close relatives) was detected in *I. multifiliis* (Zaila et al., 2017), particularly when experimentally fed on RMS-affected trout (Pasqualetti et al., 2021). Additionally, *I. multifiliis* has also been detected during trout cohabitation experiments aimed to test RMS transmission (von Gersdorff Jørgensen et al., 2019b).

In any case, besides its strong association with RMS lesions and occasional detection in other fish organs (Cafiso et al., 2016), the life cycle of RMS-MLO remains unknown, including its persistence in aquacultures after outbreaks and the potential role of hosts other than trout as vectors or reservoirs. To shed light on these still elusive aspects, here we aimed to investigate the persistence of RMS-MLO in an aquaculture facility located in Northern Italy, as well as the potential involvement in RMS of vectors or reservoirs, particularly *I. multifiliis*. We adopted a comprehensive approach, independent of RMS diagnosis in trout. Namely, we conducted for the first time a monthly sampling irrespective of RMS outbreaks for the course of one year, leveraging environmental DNA (eDNA) analyses, complemented by occasional convenience sampling from trout. eDNA-based approaches are non-invasive, and offer otherwise unattainable insights into biological communities (Ficetola et al., 2008), enabling the detection of the total genetic material within a sample, either released in the environment or present in live cells (Stewart, 2019). Thus, eDNA analyses are well suited and increasingly employed for monitoring and prevention of parasites and pathogens (Bass et al., 2023), particularly in aquaculture systems (Arbon et al., 2025; Bastos Gomes et al., 2017; Benedicenti et al., 2024), including one study focused on RMS outbreaks (Bruno et al., 2023). Notably, this approach allowed us to reveal that RMS-MLO can stably persist in aquaculture, and may interact with microeukaryotes present in the fish tanks, possibly including *I. multifiliis*, which might play a role in RMS outbreaks.

## 2. Materials and Methods

### 2.1 Sampling site

The sampling for this study was conducted in a trout farm in the province of Trento, Italy. The site is an open aquaculture system utilising freshwater sourced directly from a well. The water was initially conveyed to a primary collection tank without trout, labeled “0” (tank row 0; Figure 1). From this tank, the water was distributed through four parallel and independent trout-rearing tanks, separated by walls preventing water sharing, and labeled “A”, “B”, “C”, and “D” (tank row 1). The water then flowed into a subsequent set of four parallel and independent tanks which also housed trout, labeled “E”, “F”, “G”, and “H” (tank row 2; Figure 1).

**Figure 1.**
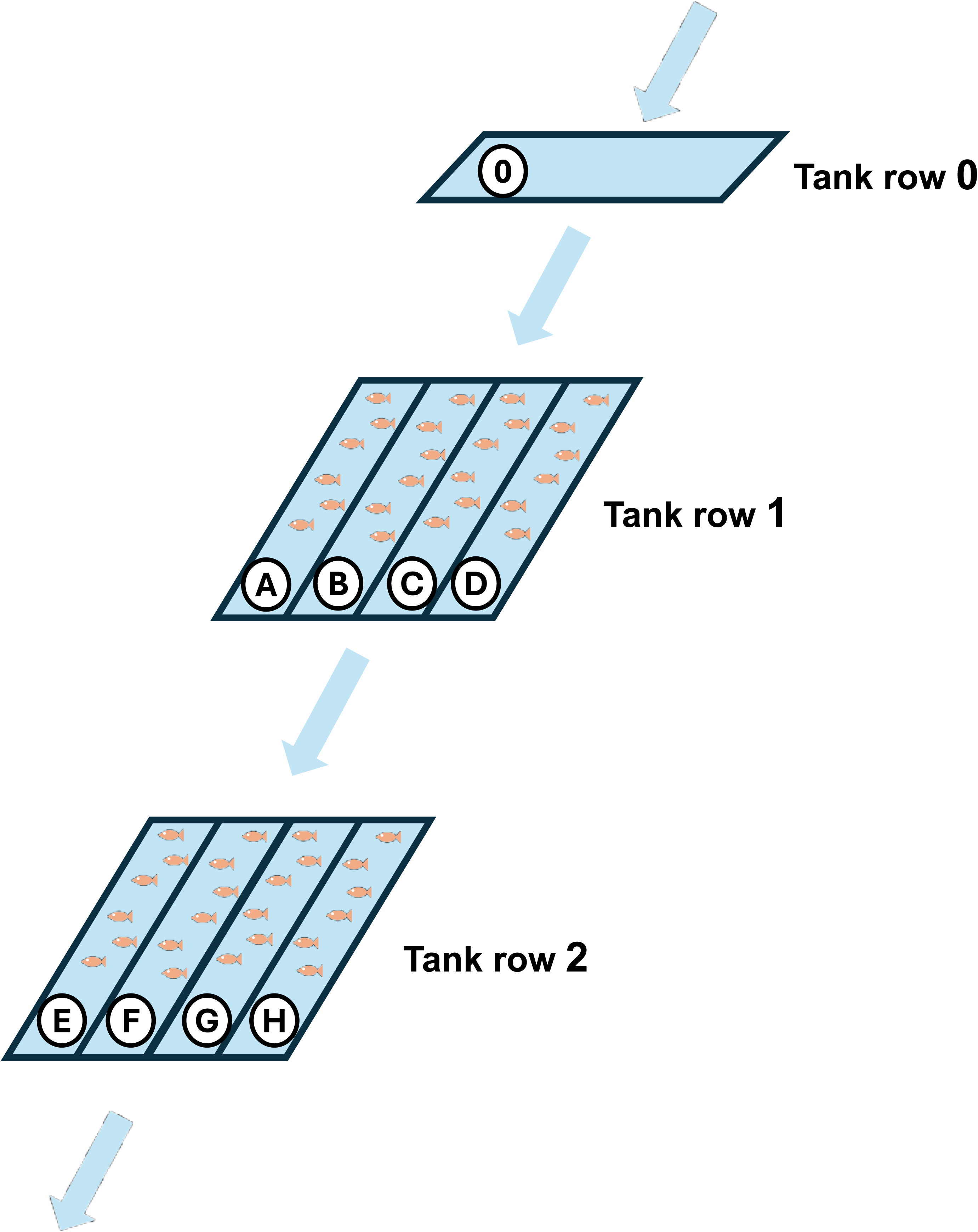
Scheme of the sampling site in the trout farm. The initial tank, labeled “0”, received fresh water from a well and contained no fish, while all the following others, labeled “A”-”H”, did. The arrows indicate the direction of water inflow, flow through the tanks, and outflow. The tank rows are indicated on the side.

### 2.2 Water and sediment sampling

Sampling was conducted over a time span of approximately one year, from 29^th^ July 2024 to 25^th^ June 2025, for a total of 11 regularly spaced timepoints.

For each timepoint, samples were taken from the nine tanks, namely the primary water collection tank, devoid of trout (“0”), and each of the eight tanks containing trout (“A”-”H”) (Figure 1). Each tank was always sampled in the same position, namely and its end point. From each tank, five litres of water were collected using five separate one-litre plastic bottles, as well as a sediment sample, by scraping the tank floor with a square plastic container. Accordingly, a total of 495 water samples and 99 sediment samples were collected throughout the study. All the reusable sampling equipment was sterilised after each use in the laboratory through UV-irradiation, or in the field using bleach, followed by rinsing with deionised water.

For each tank, we also recorded water temperature and pH (with a HI-83141-1 Analog pH/mV/temp meter; HANNA instruments), along with oxygen concentration (with an OxyGuard Handy Polaris oximeter).

During environmental sampling, the health status of the trout was visually monitored for the presence of gross skin lesions consistent with RMS (M. Galeotti et al., 2021), and the farm personnel were interviewed regarding the occurrence of RMS outbreaks or other possible infections during the previous month.

### 2.3 DNA extraction from water and sediment samples

All fresh samples were stored at 4°C, and processed within 24 h upon collection. The water from each individual sampled bottle was filtered through a separate 0.45 µm nitrocellulose filter (Thermo Fisher Scientific Inc.) using a vacuum pump. Filtration included a negative control (deionised water). Following previous studies (Cananzi et al., 2025; Morbidelli et al., 2025), filters were then cut into pieces under a sterile hood and processed using the Qiagen DNeasy® Blood & Tissue Kit. The standard protocol for tissue was followed with minor modifications, namely initial volumes of 500 µL of ATL buffer and 50 µL of proteinase K, followed by incubation at 56°C for 2 hours.

Sediment samples were initially centrifuged at 4°C for 5 minutes at 3000 *g* to remove the supernatant. Subsequently, the pellets were processed using the Qiagen DNeasy® PowerSoil® Kit, following the manufacturer’s protocol, particularly using up to 250 mg for each sample.

In all extraction sessions, a negative extraction control was included.

### 2.4 PCR analyses of water and sediment samples

All the individual environmental samples were screened by real-time PCR for the presence of RMS-MLO and *I. multifiliis*. To verify the success of the DNA extractions and the absence of PCR inhibitors, a preliminary universal bacterial PCR targeting the 16S rRNA gene was performed with Takara ExTaq and reagents (Takara Bio, Japan), the primers S-D-Bact-0008-d-S-20 (Klindworth et al., 2013) and 1492R (Matsuo et al., 2021), applying the following thermal protocol: 94°C 3’; followed by 35 cycles (94°C 30”, 53°C 30”, 72°C 2’), and 72°C 5’. Negative samples were diluted 1:10 and re-tested until a positive amplification was achieved.

To detect the presence of RMS-MLO a real-time PCR assay targeting its 16S rRNA gene was performed using primers (16SrDNA-F: 5’-GCGGTTATCTGGGCAGTC-3’; 16SrDNA-R: 5’-TGCGACACCGAAACCTAAG-3’) and protocol by (Cafiso et al., 2016), with the modifications introduced by (von Gersdorff Jørgensen et al., 2019a).

To detect the presence of *I. multifiliis*, a second real-time PCR assay targeting its 18S rRNA gene was applied, using specific primers (IMRf1: 5’-AGTGACAAGAAATAGCAAGCCAGGAG-3’; IMRr1: 5’-ACCCAGCTAAATAGGCAGAAGTTCAA-3’) (Jousson et al., 2005) with the following protocol: 94°C 3’; followed by 35 cycles (94°C 10”, 58°C 30”, 72°C 30’’), and a standard melt curve from 55°C to 95°C.

All real-time PCR reactions were conducted with Biorad reagents on a Biorad CFX Connect real-time PCR machine, testing each sample in triplicate. As a control, the amplification of a subset of samples for each assay was confirmed via agarose gel electrophoresis and Sanger sequencing (Eurofins Genomics, Germany).

### 2.5 Collection and processing of samples from trout

When possible, namely at several timepoints, additional samples were collected from individual trout. Trout collection was performed by the farm personnel as a part of their routine management. Immediately after capture, fish were externally inspected for the presence of skin lesions that could be linked to RMS. Both healthy and RMS-affected fish were then dissected for collecting samples, including diseased and healthy skin, and biopsies of multiple organs (gills, heart, liver, kidney, spleen, gut, brain, and gonads). Moreover, to increase the detection sensitivity, whole-skin surface swabs of healthy and diseased individuals were also performed. All samples were stored at -20°C. DNA extraction was performed using the Qiagen DNeasy® Blood & Tissue Kit, and real-time PCR assays for the presence of RMS-MLO and *I. multifiliis* were carried out as described above.

### 2.6 Collection and processing of samples of other potential hosts

Further water samples were collected from algae-covered sidewalls of the tanks, by scraping their surfaces with 50 mL tubes, with the aim of capturing small eukaryotic organisms that may represent additional reservoir hosts or vectors for RMS-MLO. Fresh samples were observed using a SMZ745T Nikon (Japan) stereomicroscope. Protists and small metazoans were preliminarily identified using standard taxonomic keys, manually isolated with glass micropipettes, and fixed directly in ethanol or, when possible, following laboratory propagation. Culturing was mainly achieved for ciliated protists, which were maintained in an incubator under controlled conditions (18–19°C; 12 h light–12 h dark photoperiod) and fed with *Dunaliella tertiolecta* and/or *Raoultella planticola*.

DNA extraction was performed using the NucleoSpin™ Plant II kit (Macherey-Nagel™, Germany) for protists and micrometric metazoans, and the NucleoSpin Tissue kit (Macherey-Nagel™, Germany) for other metazoans. Given the relatively low input (individual metazoans or up to ten protist cells), the quality of the DNA extraction was preliminary tested by amplification of the 18S rRNA or COI genes. The following almost universal primer pairs were used: 18S F9 Euk (Medlin et al., 1988) and R1513 Hypo (Petroni et al., 2002), LCO1490 and HCO2198 (Folmer et al., 1994), respectively. For 18S rRNA amplification in taxa such as gastrotrichs and tardigrades, group-specific primers were alternatively employed: S30 and 1806R (Norén and Jondelius, 1999); and SSU_F04 (Banerji et al., 2018) and 18S 9R (Giribet et al., 1996), respectively.

The following thermal protocols were used, namely for 18S gene: 35 cycles (94°C 30”, 54°C 30”, 72 °C 2’), and for COI gene: 40 cycles (94°C 40”, 48°C 1’, 72°C 1’). Positive samples were subsequently screened for the presence of RMS-MLO by real-time PCR as described above.

Based on PCR screening results, additional freshly isolated samples were processed for fluorescence *in situ* hybridisation (FISH), according to a previously established protocol (Nitla et al., 2019). Hybridisation experiments were performed following (Manz et al., 1992), using 30% formamide to increase probe specificity. Bacterial cells were detected using the nearly universal probe EUB338 (Amann et al., 1996), labelled with fluorescein isothiocyanate (FITC), in combination with the newly designed RMS-MLO-specific probe RMS-MLO_169 (5′-CTCGGCAATATGCAATATTAG-3′), labelled with Cyanine 3 (Cy3). The specificity of this probe was assessed by TestProbe on the SSU-rRNA SILVA r138.2 database (Chuvochina et al., 2026). Slides were examined using a Nikon microscope (Nikon Microscopy, Tokyo, Japan) equipped with a DS-Fi3 digital camera and operated with NIS-Elements D v.5.41.00 software.

### 2.7 Statistical analyses

To assess whether RMS-MLO and *I. multifiliis* exhibited temporal and/or spatial patterns, and to explore the factors potentially influencing their occurrence and co-occurrence, we applied a generalized linear mixed modelling (GLMM) framework. The presence of RMS-MLO in water samples was analysed as the number of positive samples per “batch” of samples taken from the same tank at the same timepoint (from 0 to 5), using GLMM with a Poisson distribution, including the tank as a random effect. The fixed effects considered were water tank row, sampling timepoint, number of samples positive for *I. multifiliis* in the respective batch, presence of trout, pH, temperature, oxygen concentration, oxygen saturation, and presence of a RMS outbreak. The most important predictors in affecting RMS-MLO were assessed through dredge model selection, performed by means of Akaike’s Information Criterion (AIC). The significance of each individual predictor included in the selected model was evaluated with a type II Wald chi-square test, as well as with GLMM fit by maximum likelihood with the Laplace approximation (glmerMod function). The same analyses were performed for *I. multifiliis*, replacing the fixed term “number of *I. multifiliis* positive samples in the respective batch” with the “number of RMS-MLO positive samples in the respective batch”. Prior to model fitting, multicollinearity among fixed effects was assessed (using variance inflation factors and correlation matrices), and highly collinear predictors were excluded from the analyses. To ensure the statistical validity of the final models, model assumptions were evaluated by inspecting residual diagnostics and testing for overdispersion, which allowed to verify the adequacy of the Poisson error distribution and overall model fit. All analyses were conducted using R version 4.4.2 (2024-10-31), with packages *lme4* (Bates et al., 2015), *epitrix* (Jombart et al., 2025), *performance* (Lüdecke et al., 2021), *MuMIn* (Bartoń, 2025) for statistical analysis and *dplyr* (Wickham et al., 2026), *tidyverse* (Wickham et al., 2019) and for data management and visualisation.

## 3. Results

### 3.1 Field and laboratory analyses of the environmental samples

During this study, we conducted a one year-long sampling in a rainbow trout aquaculture facility to monitor the presence and abundance of RMS-MLO and of its putative reservoir/vector *I. multifiliis*. At 11 evenly spaced timepoints, we took five water samples and one sediment sample were collected from each of the nine tanks, always in the same positions, for a total of 495 water and 99 sediment samples. A RMS outbreak in trout was detected only once (timepoint IX; April 2025). Overall, 36.6% water samples (181/495) and 53.5% sediment samples (53/99) tested positive for RMS-MLO (Figure 2a,b). When considering the five water samples jointly taken from the same tank at each timepoint as a joint batch, at least one sample was positive in the majority of batches (67.7%; 67/99; Figure 2a). A strict correspondence between water and sediment was not observed, with several instances in which a tank at the same timepoint was at least partly positive in one case and fully negative in the other, or vice versa. A more detailed examination of the data sharply revealed that the trout-free primary collection tank (tank 0) was always consistently negative across all sampling timepoints in both water and sediment. In contrast, the other eight trout-containing tanks showed considerable variations in their positivity to RMS-MLO, with the number of timepoints with at least one positive sample ranging from two (tank B, both in water and sediment), to all eleven (tank E, in water only). Besides the lack of positives in sediment at timepoint IV, no sharply evident temporal trend was observed. In water samples, tanks with positive samples ranged from four (timepoint IV) to all eight (timepoint VI) in water, while in sediment there were up to seven positive tanks (timepoints X and XI). Notably, the recorded RMS outbreak event (timepoint IX) did not correspond to a markedly higher detection of RMS-MLO in water or sediment with respect to other timepoints.

*I. multifiliis* was detected much less often, with 10.9% water samples (54/495) testing positive, corresponding to 30.3% (30/99) batches of samples taken from the same tank at the same timepoint with at least one positive sample, and 8.1% positive sediment samples (8/99) (Figure 2c,d). As observed for RMS-MLO, no strict correspondence was observed between water and sediment, nor any obvious correspondence with the respective positivity to RMS-MLO in the same sample. Interestingly, the highest presence of *I. multifiliis* in both water and sediment was observed in correspondence to the RMS outbreak (timepoint IX) and in the subsequent timepoint X. This timepoint also involved a relatively high number of positive water samples (3 and 4, respectively) in the trout-free primary collection tank (tank 0) (Figure 2c).

**Figure 2.**
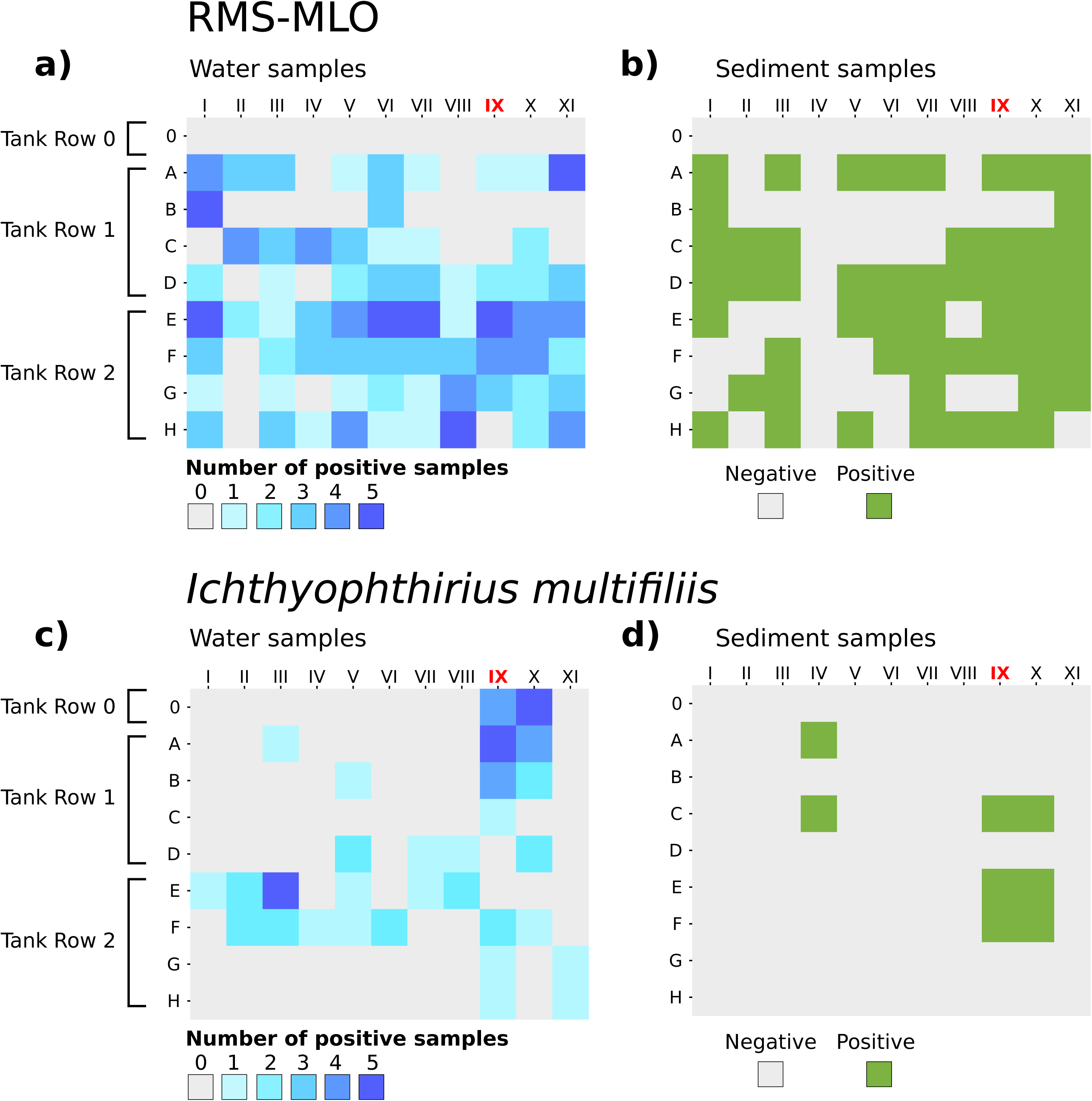
Heatmaps indicating the positivity to RMS-MLO in water (a) and sediment (b) samples, as well as to *I. multifiliis* in the same samples (c,d). In (a,c), the number of positive water samples (from zero to five) per tank and timepoint is shown in different shades of blue, while in (b,d) the positivity of each single sediment sample taken is shown in green. The timepoint at which a RMS outbreak was detected (IX) is shown in red. Image realised with chiplot.online (https://www.chiplot.online/).

Among the recorded environmental parameters, pH (range 5.87-7.76) and oxygen levels (range 5.01-15 mg/L) showed relatively large temporal and spatial variations, while temperature (range 10.7-11.7°C) was more stable, with its highest values at the timepoints I-III, corresponding to the summer 2024 (Supplementary Figure 1).

### 3.2 Statistical analyses on the presence of RMS-MLO and *I. multifiliis* in water samples

Next, we aimed to analyse the influence of different factors on the presence of RMS-MLO and *I. multifiliis* in the water samples. The most parsimonious model describing the number of positive samples of RMS-MLO in each batch included tank row as a fixed effect and tank as a random intercept. Model diagnostics did not show overdispersion nor singularity of the random effects structure, thus indicating an overall adequacy of the model. The effect of tank row suggested an increasing trend in the number of positive RMS-MLO samples along the water flow, however, this was not statistically significant (Wald χ²=4.84, df=2, p=0.089). Among the other variables examined, the trout presence was excluded from modeling due to its complete collinearity with tank row, as trout were always absent in row 0 and always present in rows 1 and 2. On the other hand, while the most parsimonious model did not include any of the other variables, particularly RMS outbreak and *I. multifiliis*, as well as environmental abiotic factors, the relationship of those variables with RMS-MLO abundance cannot be excluded based on this dataset, as the collinearity structure did not allow a fully reliable assessment.

In contrast, the most parsimonious model describing the number of positive samples per batch to *I. multifiliis* was included tank as random intercept and, as fixed effects, outbreak occurrence, pH and number of RMS-MLO positive samples, without multicollinearity among the predictors. Moreover, the overall adequacy of the model was confirmed as well by the lack of overdispersion and singularity. A significant positive effect of outbreak occurrence (p<0.001) and significant negative effects of RMS-MLO relative abundance (p<0.05) and pH (p<0.01) were detected. These results indicate an increased probability of detecting *I. multifiliis* during RMS outbreaks, and a decreased probability of detection at higher RMS-MLO abundance and under more alkaline pH conditions.

### 3.3 Molecular screening for RMS-MLO and *I. multifiliis* in trout

Given the high and relatively constant presence of RMS-MLO in water and sediment samples throughout the study, particularly in the absence of overt RMS outbreaks, we also investigated the presence of this bacterium directly in fish. Therefore, convenience samplings from trout were performed. A total of 28 samples were collected at timepoint VIII, which preceded the RMS outbreak, from 10 healthy trout (10 whole skin swabs, 10 gill biopsies, and 8 internal organ biopsies) (Supplementary table S1). All samples were negative in PCR for both RMS-MLO and *I. multifiliis*. During the RMS outbreak (timepoint IX), 27 samples were taken from nine RMS-affected trout, in most of which (66.7%) RMS-MLO was molecularly detected (9/9 skin lesions, 7/9 whole skin swabs, and 2/9 gill biopsies). In addition, *I. multifiliis* was detected in one whole skin swab and one skin lesion sample, both also positive to RMS-MLO (Supplementary table S1). Additional fish samples were collected at timepoint XI, including from three fish individuals showing RMS lesions (although in the absence of an overt outbreak), two fish with skin lesions not attributable to RMS, and five fish without lesions. Out of 25 samples (10 whole skin swabs, five skin lesion samples, and 10 gill biopsies), RMS-MLO was molecularly detected in skin lesions from the three RMS-affected fish, while all other samples, including those from fish with non-RMS lesions, tested negative. None of these samples tested positive for *I. multifiliis*.

### 3.4 Detection of RMS-MLO in other hosts

To gather insights on the potential association of RMS-MLO with other aquatic eukaryotic organisms (e.g., protists, small metazoans) and their possible role as reservoirs or vectors, further water samples were collected at multiple timepoints and inspected at the microscope. Several diverse organisms were isolated, ranging from arthropods, rotifers, tardigrades, flatworms, annelids, ciliates, and amoebas (Supplementary table S2). Real-time PCR screening for the presence of RMS-MLO yielded an appreciable amount of positive samples (7.0%; 5/7), spanning multiple taxa, including ciliates (*Anteholostica monilata*, *Apodileptus vischeri rhabdoplites*), testate amoebae (*Centropyxis* sp.), oligochaete annelids (*Chaetogaster* sp.), and an isopod. Therefore, in order to investigate possible direct interaction between RMS-MLO cells and potential hosts other than trout, further newly isolated individuals of free-living microeukaryotes were inspected by FISH. To this end, the RMS-MLO_169 probe, specific for RMS-MLO, was designed on purpose (no off-target hits allowing up to one mismatch, only six allowing two mismatches), and used in combination with an almost universal bacterial probe as control. Accordingly, we detected positive probe signals consistent with RMS-MLO cells within food vacuoles of multiple ciliates, namely euplotids, prostomateans and a *Paramecium* sp. (Supplementary Fig. S2).

## 4. Discussion

The RMS is a worldwide distributed skin disease affecting farmed rainbow trout, with significant economic impact (Ferguson et al., 2006; Orioles et al., 2023; Schmidt et al., 2018). Despite its relevance, its aetiology and epidemiology are still poorly understood (Metselaar et al., 2022). The putative causative agent is a *Rickettsiales* bacterium, termed RMS-MLO, which is however known almost exclusively from PCR-based detection from trout skin lesions (Cafiso et al., 2016; Lloyd et al., 2011; Metselaar et al., 2010), plus, in a few instances, from putative electron micrographs (Galeotti et al., 2017a; Orioles et al., 2022a). These data derived from diseased trout sampled during overt RMS outbreaks in trout farms, but, to our best knowledge, no data exist on the occurrence and dynamics of the bacterium outside such events, also considering the limited data on their exact frequency and duration (Metselaar et al., 2022; von Gersdorff Jørgensen et al., 2019a).

In this work, we aimed to fill this gap of knowledge by focusing on the analysis of eDNA, an innovative approach, unbiased with respect to RMS diagnosis in trout. eDNA analyses are suitable for epidemiological investigations and long-term monitoring given their inherent non-invasive nature. Specifically, they are particularly convenient in aquaculture settings, since they do not impact fish nor interfere with routine farming activities. This allowed us a continuous sampling over an entire year, providing an unprecedented temporal resolution for assessing the dynamics of RMS-MLO and its relation to RMS outbreaks. While innovative and yet not fully exploited, eDNA-based approaches are now well established, and have been widely used for studying the distribution and abundance of aquatic organisms (Ficetola et al., 2008; Stewart, 2019), including parasites and pathogenic microorganisms (Arbon et al., 2025; Bass et al., 2023; Bastos Gomes et al., 2017). Particularly, RMS-MLO well exemplifies the technical establishment but still underexploitance of eDNA analyses, considering that, although an eDNA-targeted molecular assay was already successfully tested and applied, its use has been so far limited to RMS outbreaks (Bruno et al., 2023).

Our main finding was that, contrary to expectations, RMS-MLO was consistently and abundantly detected throughout the year, without any appreciable temporal variations, nor, chiefly, any significant increase linked to the single RMS outbreak recorded during this study. It is noteworthy that, while often detected in each of the tanks housing trout (rows 1 and 2), this bacterium was always fully absent in the primary collection tank (row 0), which contained no fish and was supplied exclusively by incoming water (Figure 2a,b). This suggests that, at least during the study period, this bacterium was not introduced via the inflow water, but rather was already present and able to persist at relatively high abundance within the trout tanks. Moreover, our statistical analysis showed a potential, though non-significative, increase of RMS-MLO according to the tank row. This trend may suggest a potential accumulation of the bacterium along the tanks (Figure 2a,b), but further targeted analyses will be required to confirm this result. To our best knowledge, this is the first report of RMS-MLO in a rainbow trout aquaculture setting in the absence of overt RMS outbreaks, as well as of its persistence throughout all seasons over a full year. These observations could support previous hypotheses on the geographical spread of RMS due to the exchange of trout, eggs or water from farm to farm (Metselaar et al., 2022), rather than originating from upstream water sources, thus highlighting the need for targeted preventive actions.

The lack of a clear temporal link between environmental RMS-MLO detection and RMS outbreaks could hypothetically be the result of residual eDNA of this bacterium as a consequence of previous outbreaks. However, this scenario seems highly unlikely, considering that the investigated timespan of one year largely exceeds known eDNA persistence (few days/weeks) (Barnes et al., 2014; Jo and Minamoto, 2021; Lamb et al., 2022), and the regular cleaning procedures in the tanks operated by the aquaculture personnel (personal communication). Moreover, the possibility of a long-term persistence of RMS-MLO in a free-living stage is also very improbable, considering the known and ancient obligate host-association of the *Rickettsiales* (Castelli et al., 2024; Salje, 2021), as well as previously failed cultivation attempts of this bacterium (Metselaar et al., 2022). These observations strongly indicate that RMS-MLO persisted during our study in a host-associated stage, for which multiple non-mutually exclusive possibilities can be considered. The most obvious hypothesis on the potential host of RMS-MLO between outbreaks would be the trout themselves, consistent with the strong association observed between this bacterium and trout presence, in contrast to its absence in the trout-free inflow tank. This would imply a somehow opportunistic behaviour of RMS-MLO, and/or the presence of RMS cases at low frequency, below the detection thresholds during non-invasive and routine observation of the bulk fish mass. Considering the detection of a small number of trout with RMS-MLO-positive skin lesions outside the outbreak period (Supplementary Table S1), our data support the latter hypothesis, and align with previous reports regarding the negativity of healthy fish (Cafiso et al., 2016). Further in-depth analyses, inevitably more invasive, would be necessary in the future to clarify this point, as well as to assess any potential quantitative relationship between RMS prevalence in trout and environmental detection of RMS-MLO.

Previous studies indicated an association between RMS-MLO and *I. multifiliis* (Pasqualetti et al., 2021; Zaila et al., 2017). However, our in-depth data argue against a relevant reservoir role of this parasite, considering its much lower occurrence as compared to RMS-MLO (Figure 2) and the negative statistical association found between the two. On the other hand, we found indications of at least transient interactions between RMS-MLO and free-living microeukaryotes, particularly other ciliates (Supplementary Table S2; Supplementary Figure S2). While the ascertained presence of this bacterium only in food vacuoles does not confirm a stable interaction, this condition could anyway represent the basis also for long-term persistence (Modeo et al., 2020), thus leaving open the possibility of at least occasional reservoir hosts for RMS-MLO, also accounting for the wide spectrum of host adaptations of the *Rickettsiales* among unicellular and multicellular aquatic organisms (Castelli et al., 2024, 2016).

Despite the overall rarity of *I. multifiliis* in our samples (Figure 2c,d), which seems in apparent disagreement with previous studies on its role in RMS (Pasqualetti et al., 2021; Zaila et al., 2017), we found a statistically supported link between its relatively much higher abundance in a few timepoints (IX and X) and the concomitant occurrence of the RMS outbreak. This hints for a relevant involvement of this parasite in RMS, possibly as a facilitator, consistent with the higher presence of RMS-MLO-like bacteria in other fish species that are susceptible to *I. multifiliis* (Liu et al., 2025). For instance, it could favour the colonisation of fish skin by physically transporting RMS-MLO through a transient interaction (Pasqualetti et al., 2021; Zaila et al., 2017), or as a consequence of some synergistic action (Okon et al., 2023), such as inducing physical damages on the fish (Wang et al., 2019). Particularly, the high abundance of *I. multifiliis* recorded in the primary collection tank (tank 0, Figure 2c,d) suggests that it could be introduced from external sources, in contrast to RMS-MLO. These findings may imply that *I. multifiliis* could act a trigger for RMS outbreaks in aquacultures where RMS-MLO is already present. Therefore, future RMS management strategies could account for the control of *I. multifiliis*, despite it not being the primary aetiological agent.

While previous studies suggested a role of seasonal cycles in RMS, due to its association with cold temperatures (Metselaar et al., 2022; Oidtmann et al., 2013; Orioles et al., 2022a), here we recorded a single outbreak in spring (timepoint IX: April 2025). Moreover, we detected very modest temperature variations throughout the study (Supplementary Figure S1), which, unsurprisingly, were not linked to any relevant effect on RMS-MLO (nor on *I. multifiliis*). Nevertheless, other environmental parameters were more variable (Supplementary Figure S1), particularly a more acidic pH was significantly associated with a higher presence of *I. multifiliis*, consistent with previous records of pH modulating its reproductive cycle (Garcia et al., 2011; Tange et al., 2020). Given the possible involvement of *I. multifiliis* in RMS, targeted experiments on the effect of pH on disease transmission, possibly based on trout cohabitation assays (Orioles et al., 2022a), may help elucidate its role in shaping the interactions between host, pathogen and environment, as well as the disease onset.

Taken together, our findings highlight the complexity of RMS epidemiology in aquaculture systems and the advantages of eDNA analyses for its investigation and monitoring. Future research will be necessary to clarify the relationship between the environmental abundance of RMS-MLO and disease incidence, the involvement of *I. multifiliis* and its actual interactions with RMS-MLO, as well as the role of trout physiology (e.g., previous exposure to RMS-MLO) in the disease onset. Moreover, a better understanding of the ecological dynamics between RMS-MLO and other organisms inhabiting aquaculture systems, including potential reservoirs, may be relevant to unravel the processes underlying disease outbreaks. Addressing these knowledge gaps will not only be fundamental to clarify the aetiology of RMS, but will also allow the development and implementation of targeted and sustainable disease management strategies in rainbow trout aquaculture. This may ultimately shift the focus from the pathogen itself towards the control of ecological and environmental factors that lead to the emergence of outbreaks.

## Supporting information

Legend of the Supplementary material

## Acknowledgements

This work was funded by the Italian Research Ministry PRIN project n. 2022283TH9 to MC, PT, and VS. Gabriele Cananzi and Irene Tatini are gratefully acknowledged for helpful suggestions on protocols for sampling and laboratory analysis of eDNA. The authors are grateful to the staff of the aquaculture facility involved in this study for their valuable cooperation, availability, and assistance with sampling activities throughout the monitoring period.

