## Supplementary material for "Environmental DNA reveals long-term persistence of a *Midichloria*-like bacterium in a rainbow trout aquaculture and links *Ichthyopthirius multifiliis* with the red mark syndrome": Legend of the Supplementary material

**Legend of the Supplementary Materials**

**Supplementary Figure S1.** Heatmaps indicating pH, temperature and oxygen concentration values recorded across the samplings in coloured scales from yellow (lowest) to red (highest). The timepoint at which a RMS outbreak was detected (IX) is shown in red. Image realised with chiplot.online (https://www.chiplot.online/).

**Supplementary Figure S2.** Fluorescence *in situ* hybridization experiments on ciliates. (A–C) The same euplotid cell showing the macronucleus (MAC) stained with DAPI (A), a positive signal from the universal bacterial probe EUB338 in food vacuoles (arrows), emitting green fluorescence (B), and a positive signal from the species-specific probe for RMS-MLO, Red_Mark_169, in some food vacuoles, emitting red fluorescence (arrowheads) (C). (D–F) The same prostomatean cell showing the macronucleus (MAC) stained with DAPI (D), a positive signal from the universal bacterial probe EUB338 in food vacuoles (arrows), emitting green fluorescence (E), and a positive signal from the species-specific probe for the RMS-MLO, Red_Mark_169, in some food vacuoles, emitting red fluorescence (arrowheads) (F). (G–I) The same two *Paramecium* cells showing the macronucleus (MAC) stained with DAPI (G), a positive signal from the universal bacterial probe EUB338 in food vacuoles (arrows), emitting green fluorescence (H), and a positive signal from the species-specific probe for the RMS-MLO, Red_Mark_169, in some food vacuoles, emitting red fluorescence (arrowheads) (I). MAC, macronucleus; arrows indicate food vacuoles; arrowheads indicate bacteria positive for the probe Red_Mark_169. Scale bars: 10 µm (A–F) and 50 µm (G–I).

**Supplementary Table S1.** Summary of the results of the real-time PCR screening for the presence of RMS-MLO and *Ichthyphthirius multifiliis* in trout samples. Samples are arranged by sampling timepoint (Roman numerals), while samples with the “trout ID” were collected from the same fish individual.

**Supplementary Table S2.** Summary of the results of the real-time PCR screening for the presence of RMS-MLO in samples from protists and small metazoans. Samples are arranged according to respective taxonomic groups.
