## Supplementary figures and images for "Environmental DNA reveals long-term persistence of a *Midichloria*-like bacterium in a rainbow trout aquaculture and links *Ichthyopthirius multifiliis* with the red mark syndrome"

### Supplemental Figure 1

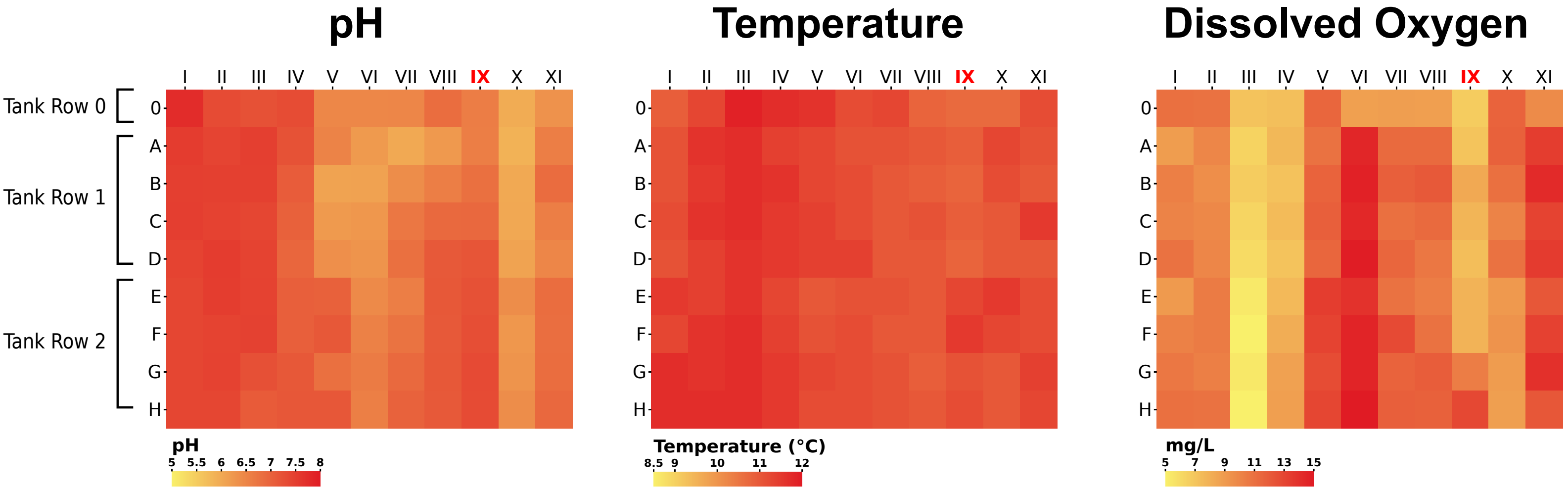

### Supplemental Figure 2

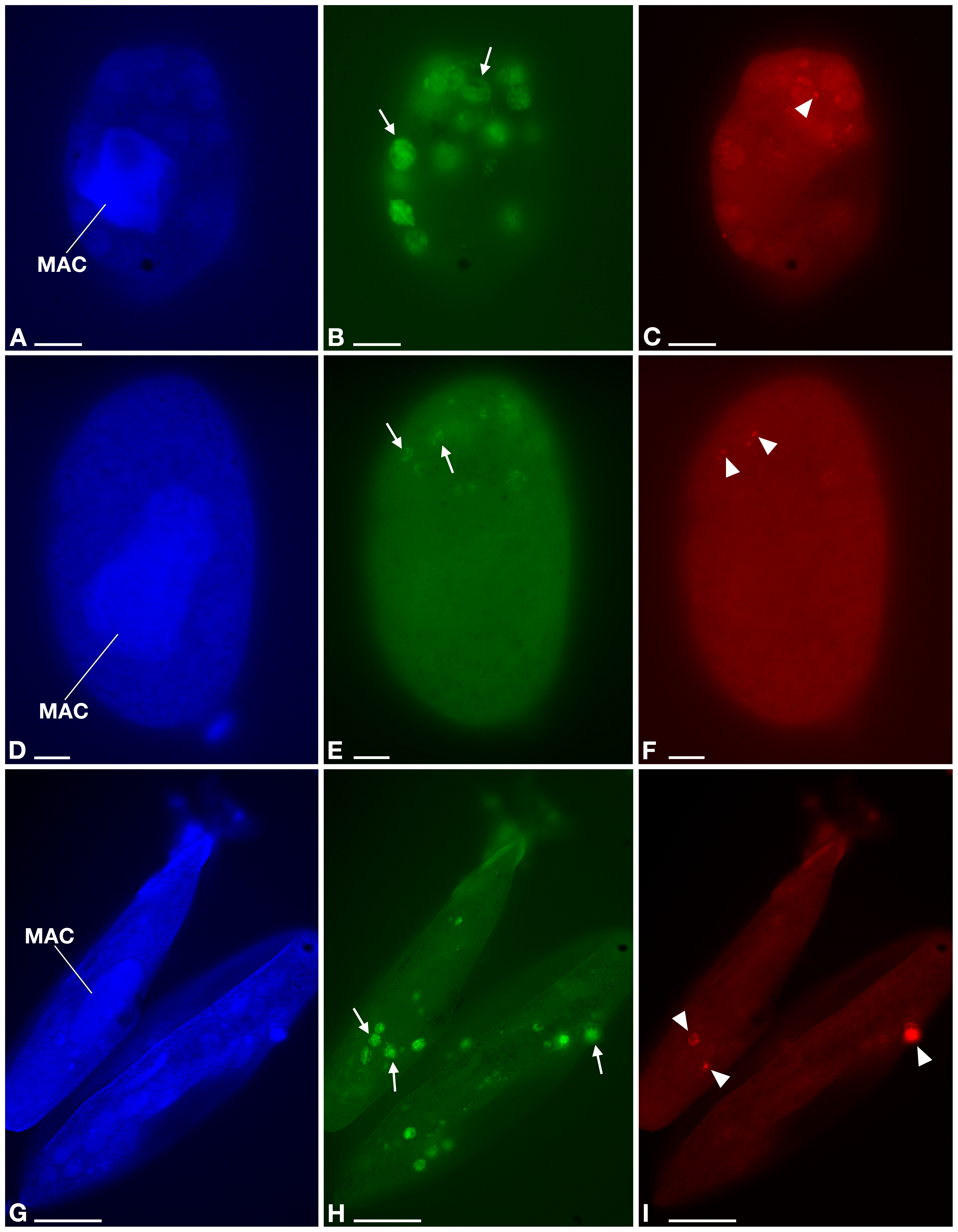
